# Global Profiling of N-terminal Cysteine-Dependent Degradation Mechanisms

**DOI:** 10.1101/2025.01.20.633921

**Authors:** Aizat Bekturova, Yaara Makaros, Shahar Ben-David, Itay Koren

## Abstract

Hypoxia, a condition characterized by insufficient oxygen supply, challenges cellular homeostasis and energy production, driving the activation of adaptive responses to maintain survival under these stress-inducing conditions. One key strategy involves enzymatic oxidation of N-terminal cysteine residues coupled with proteolysis through the Cys-Arg/N-degron pathway. Despite the presence of hundreds of proteins with N-terminal cysteine in humans, only two have been identified as substrates of this pathway, and its substrate selectivity remains unclear. Moreover, the biological role of this pathway in the cellular response to hypoxia is not well defined. By employing a systematic proteomics approach, we discovered that nearly half of the cysteine-commencing proteome could be regulated by the Cys-Arg/N-degron pathway. Mutagenesis experiments revealed the specificty of Cys-Arg/N-degron pathway showing a preference for hydrophobic and positively charged residues following cysteine. Furthermore, we uncovered substrates that are regulated by this pathway during hypoxia, including IP6K1. The loss of IP6K1 impaired glucose uptake, glycolytic ATP production, and overall mitochondrial morphology and function. As a result, IP6K1-deficient cells exhibited disrupted metabolic adaptation under hypoxic conditions and decreased survival under stress. These findings underscore the importance of the Cys-Arg/N-degron pathway in regulating metabolic responses and highlight its potential importance in hypoxia-related disorders.

## Introduction

Hypoxia refers to conditions of reduced oxygen availability, which presents a major challenge to cellular function, as oxygen is essential for energy production and maintaining cellular homeostasis. In response, cells activate a range of adaptive mechanisms to adjust to oxygen levels, enabling them to manage metabolic stress, preserve function, and survive under these oxygen-deprived conditions (1). Two key regulatory mechanisms implicated in hypoxic stress response are the hypoxia-inducible factor 1 alpha (HIF-1α) pathway and the Cys-Arg/N-degron pathway of protein degradation (1–3). Among these, the HIF-1α pathway has been more extensively studied. Specifically, HIF-1α regulates the expression of genes involved in cellular adaptation to low oxygen levels, including metabolism, angiogenesis, and erythropoiesis (4).

N-degron pathways refer collectively to proteolytic pathways that selectively promote the degradation of proteins containing specific “destabilizing” N-terminal degradation motifs, known as N-degrons, to maintain cellular proteostasis and control cellular signaling (5–9). There are several types of N-degron pathways, some of which involve modification to the N-terminal residue such as the Arg/N-degron pathway, the Ac/N-degron pathway and the fMet/N-degron pathway that involve Nt-arginylation, Nt-acetylation, and Nt-formylation, respectively (6). Among N-degron pathways that regulate the turnover of proteins bearing non-modified N-termini are the recently discovered Pro/N-degron pathway (10) and GASTC/N-degron pathway (GASTC = Gly, Ala, Ser, Thr, Cys) (11–13).

Among the various N-degron pathways, Arg/N-degron pathway has been shown to function as an oxygen sensor regulating the turnover of a subset of proteins with cysteine at the N-terminus, following the initiator methionine (iMet) (hereafter refers as Cys-Arg/N-degron pathway) (14). This pathway involves a series of processing and modification steps at the N-terminus of protein substrates. First, methionine aminopeptidases (MetAPs) co-translationally remove the iMet from nascent proteins when the residue following iMet is small (e.g., Gly/Ala/Val/Ser/Thr/Pro/Cys) (15). This process exposes cysteine as the new N-terminal residue. Under normoxic conditions, the N-terminal cysteine undergoes deoxygenation by 2-aminoethanethiol dioxygenase (ADO) (2), which activates it for the next step-N-terminal arginylation. This modification is catalyzed by arginyltransferase ATE1, which adds arginine to the N-terminal oxidized cysteine, transforming the protein into a Cys-Arg/N-degron substrate. The modified protein is recognized by E3 ligases containing UBR-box domains, which facilitate its ubiquitination and subsequent degradation via the proteasome (14). This process ensures that substrates of the Cys-Arg/N-degron pathway are degraded under normoxic conditions when they are not required, but their turnover is halted under hypoxic conditions, allowing them to accumulate and possibly to play a role in the cellular response to low oxygen.

Which proteins are substrates of the Cys-Arg/N-degron pathway? In higher plants VII ETHYLENE RESPONSE FACTOR (ERFVII), a regulator of hypoxia-induced transcriptional reprogramming, is a direct substrate of the Cys-Arg/N-degron pathway (16). In humans, however, the pathway remains understudied, with the only known substrates being the GTPase-activating proteins RGS4, RGS5, and RGS16 (17) and interleukin-32 (IL32) (2). RGS proteins accumulation in ATE1^-/-^ embryos disrupts Gα_q_-mediated cardiac signaling pathways, leading to impaired cardiac function (17–19), suggesting that the Cys-Arg/N-degron pathway plays a crucial role in maintaining low levels of these proteins under normoxic conditions. However, the importance of preserving their degradation under hypoxic conditions is still unclear. A recent study has proposed that RGS5 might have a specific role in hypoxia, where it inhibits pericyte recruitment to blood vessels (20). As for IL32, it has both pro and anti-inflammatory roles and and its elevated expression has been linked increased cell proliferation and cancer progression (21). However, the biological significance of its regulation by Cys-Arg/N-degron pathways is yet to be fully understood.

Met-Cys (MC)-starting proteins comprise approximately 1% of the human protein-coding genome (279 proteins), suggesting the possibility of additional substrates regulated by the Cys-Arg/N-degron pathway. However, to date, no systematic studies have successfully identified new substrates for this pathway. Proteomic analyses following ADO depletion in cancer cells did not reveal significant changes in the proteome (22). RNA sequencing (RNA-seq) and mass spectrometry (MS) analysis conducted in human UBR1/2 knockout (KO) cells revealed significant alterations in the expression of approximately 300 genes, however this approach did not directly identify MC-initiating substrates of the Arg/N-degron pathway (23). In addition, a recent study profiling *in vitro* 68 human MC-commencing peptides to screen for potential ADO substrates found that while ADO exhibits selectivity for specific N-terminal cysteine containing motifs, no novel endogenous MC-proteins were identified (24). These limitations in sensitivity and throughput have hindered the discovery of additional protein substrates for this pathway.

To address these limitations and to identify the full scope of Cys-Arg/N-degron pathway substrates, we utilized here a global stability profiling (GPS)-peptidome approach, a high-throughput method designed to characterize degron motifs in human proteins (12,25). Using this approach, we systemically analyzed the stability of MC-initiating peptides in response to ablation of ATE1 using a library of peptides that encode the N-termini of all human proteins. Our results reveal that 45% of all MC-starting proteins are regulated by ATE1. Furthermore, mutagenesis library screens demonstrate the specificity of the Cys-Arg/N-degron pathway, highlighting distinct amino acid preferences downstream of the cysteine residue. Through these screens, we also identified previously unrecognized full-length protein substrates of the Cys-Arg/N-degron pathway, including Primary Cilia Formation (PIFO) and Inositol Hexakisphosphate Kinase 1 (IP6K1), prompting further investigation into the role of IP6K1 in the cellular response to hypoxia. IP6K1 is an enzyme that catalyzes the conversion of inositol hexakisphosphate (IP6) into 5-diphosphoinositol pentakisphosphate (5-PP-IP5), also known as IP7. These IPs play a critical role in cellular signaling, regulating processes such as energy metabolism, cell proliferation, and programmed cell death (26,27). Additionally, IP6K1 has been shown to regulate insulin sensitivity, promote weight loss, and improve glucose tolerance (28,29). Although IP6K1 is known to regulate metabolism, its role in hypoxia and the mechanisms controlling its expression remain unclear. In this study, we discovered that IP6K1 is subjected to degradation via the Cys-Arg/N-degron pathway. While under hypoxic conditions IP6K1 protein levels are elevated, due to inhibition of its degradation, reducing its expression through CRISPR/Cas9 knockout (KO) led to decreased proliferation under hypoxia. This reduced survival can, in part, be attributed to the impaired glucose uptake and disrupted metabolic adaptation observed in IP6K1-deficient cells. These findings offer valuable insights into the role of IP6K1 in oxygen sensing in eukaryotes.

## Results

### Identification of Cys-Arg/N-degron substrates by systematic stability analysis using N-terminome library in ATE1 ablated cells

To investigate the role of N-terminal cysteine in promoting protein instability and to determine the selectivity of substrate selection within Cys-Arg/N-degron pathway, we performed a GPS-peptidome screen using N-terminal peptide library in ATE1 knockout (KO) cells (Supplemental Figure S1A, Figure 1A). The GPS method (30) monitors protein stability in live cells by employing lentiviral expression of human peptidome libraries fused to GFP (12,25). The peptide-GFP fusion levels are normalized to a control protein, DsRed, which is expressed from the same transcript. The ratio of GFP to DsRed signals serves as an indicator of the stability of the GFP-fusion peptides (12,25). We recently adapted this approach by incorporating the ubiquitin-fusion technique (7) creating a GPS-peptidome system where the first 24 amino acids (hereafter “N24mer”) of human protein isoforms are fused between ubiquitin and GFP (“Ub-GPS”) (Figure 1A) (12). Upon expression of the constructs in human cells, proteolytic cleavage of the ubiquitin moiety by endogenous deubiquitinating enzymes lead to the exposure of the peptides at the N-terminus of GFP (Figure 1A). We introduced the library in duplicates into human embryonic kidney (HEK) 293T control or ATE KO cells and used fluorescence-activated cell sorting (FACS) to partition the population into four bins of equal size based on the stability of the peptide-GFP fusions. The stability of each fusion was then quantified with Illumina sequencing, with each peptide assigned a protein stability index (PSI) score ranging between 1 (maximally unstable) and 4 (maximally stable) according to the proportion of sequencing reads in each bin from control or ATE1 KO cells (Figure 1A, Supplementary Table 1). Of the 279 MC-starting peptides present in the library, 245 had sufficient reads for analysis. The screen revealed that 110 out of 245 MC-peptides showed stabilization (ΔPSI ≥0.3) in ATE1 KO cells (PSI difference between the ATE1 KO and control KO) (Supplementary Table 1). To determine whether the Cys-Arg/N-degron pathway has a preference beyond the N-terminal cysteine, we calculated the enrichment fold of residues at the third position (MC**<u>X</u>**-) in ATE1 substrates, comparing them to their distribution at the same position in the human proteome. The results revealed a distinctive amino acid composition in ATE1 substrates, with the most significantly enriched motifs among the stabilized MC-peptides in the ATE1 KO background were those containing hydrophobic (e.g., Phe, Val, Ile) and positively charged residues (e.g., His, Arg) following Cys (Figure 1B). Cross comparing with a previous N-terminome screen conducted in UBR1/2/4 triple KO cells (UBR KO) (12), revealed that approximately half of ATE1 candidate substrates also scored as UBR substrates (57 out of 110) exhibiting the same trend of N-terminal preference downstream of Cys (Supplemental Table 1). Notably, the median PSI of N-terminal cysteine peptides with preferred or non-preferred ATE1 motifs was significantly lower when the motif was located at the extreme N-terminus, as opposed to internal regions (Figure 1C). This suggests that N-terminal cysteine motifs, in general, promote protein instability regardless of their regulation by the Cys-Arg/N-degron pathway. However, among the MC motifs, those recognized by ATE1 exhibited significantly lower PSIs, indicating stronger instability conferred by the Cys-Arg/N-degron pathway.

**Figure 1.**
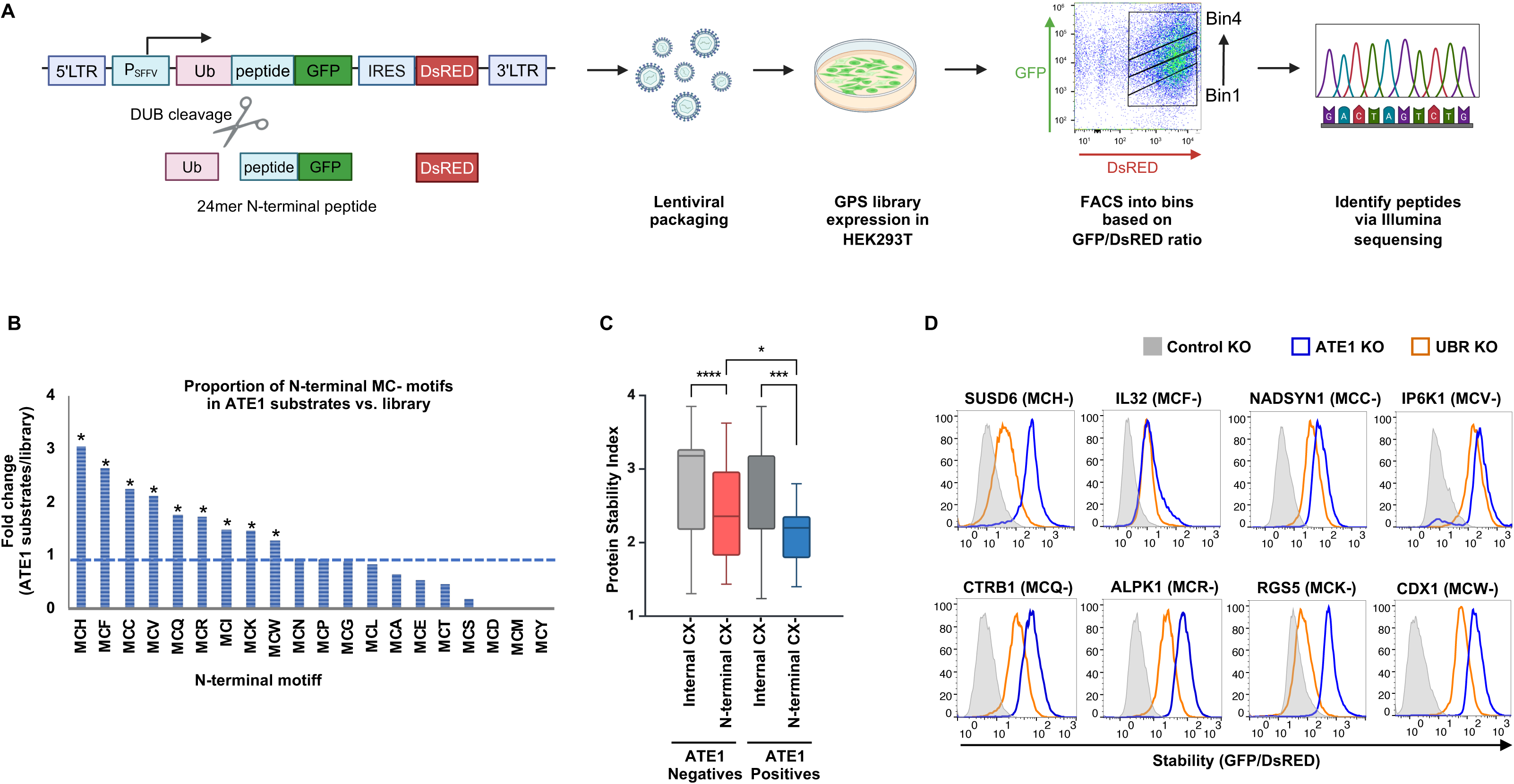
Identification of Cys-Arg/N-degron substrates through stability profiling of an N-terminome library in ATE1-deficient cells. (A) Schematic representation of the N-terminome GPS screen, in which the first 24 residues of all human proteins were expressed as N-terminal fusions to GFP in the Ub-GPS vector. To isolate peptide-GFP fusions based on their stability, fluorescence-activated cell sorting (FACS) was utilized to sort cells into four populations (“Bins”) based on their GFP/DsRed fluorescence ratio. Next-generation sequencing was used to identify the peptides enriched in each bin. Illustration was generated using BioRender. (B) The fold-change in amino acid frequency at position 3 (MC**<u>X</u>**-) of ATE1 substrates was compared to the overall frequency of reside “X” at position 3 in the library. Statistical significance for each N-terminal motif is indicated by Z-score (see Materials and Methods). (C) Boxplots showing the distribution of PSI stability scores for all peptides in which the indicated motifs were encoded at the second position (colored boxes) versus any other internal position within the peptide (gray boxes). ATE1 favored motif is presented in blue while ATE1 disfavored motif is shown in red. (D) Ub-GPS constructs, where GFP was fused C-terminally to the first 24 residues of the indicated genes were expressed in control (cells transduced with AAVS1 sgRNA), ATE1 (sg1-ATE1) or UBR1/2/4 triple KO cells (“UBR KO”). Stability was then analyzed by flow cytometry. GFP/DsRed ratio represents the stability of the indicated GFP-fusion proteins. Stabilization of the target protein in various genetic background is indicated by a sharp peak to the right side of each panel. For each gene, its name and the first N-terminal three residues are indicated.

Notably, the N-terminome screen also revealed an additional 57 non-MC starting peptides whose stability is regulated by ATE1, 18 of which were also identified in the UBR screen (Supplementary Table 1). The regulation of these substrates by both ATE1 and UBRs, despite the absence of an MC-motif, suggests that they may undergo cleavage by proteases such as dipeptidyl peptidases (DPPs) (31), calpains, separases, and secretases (32–34) which expose neo-N-termini that are recognized by the Arg/N-degron pathway. For example, CEP290, ARID1B and PCK1 are among DPP8/9 substrates (31) and contain MPPN- and MPPQ-N-terminal motifs. After cleavage, these substrates expose N- and Q-at the neo-N-terminus, which are then subjected to amidation and arginylation, converting them into Arg/N-degron substrates (31). Alternatively, some of these non-MC-peptides may undergo internal arginylation (32,35), although we cannot rule out potential indirect effects on the stability of this subset of substrates due to ATE1 KO.

To validate the results from the ATE1 screen, example candidate ATE1/UBR peptide substrates containing various N-terminal Cys motifs were cloned individually in the GPS vector, expressed in ATE1 and UBR KO cells and stability was analyzed using flow cytometry. In all cases, peptide stability was increased in ATE1 and UBR KO cells confirming the validity of the N-terminome screen and indicating that these peptides are regulated by the Cys-Arg/N-degron pathway (Figure 1D).

### Uncovering the specificity of Cys-Arg/N-degron substrate recognition through mutagenesis libraries

Next, we conducted mutagenesis experiments on a set of MC-ATE1 substrate peptides to assess the role of MC-motif in driving substrate instability. In this set of experiments, we generated peptide libraries wherein single or triple amino acid mutations were introduced across all positions within the 24mer sequence. The scanning mutagenesis libraries were cloned into Ub-GPS vector and the GPS screen of mutagenesis library was performed as described earlier. Notably, our findings revealed that mutating Cys and the adjacent 1-2 residues downstream resulted in a significant stabilization of the tested peptides (Figure 2A, Supplemental Table S2). This suggests that the motif composed of the N-terminal Cys and the immediately adjacent residues is relatively short and critical for substrate instability. Additionally, residues beyond the N-terminal fourth position do not appear to play a role in substrate turnover.

**Figure 2.**
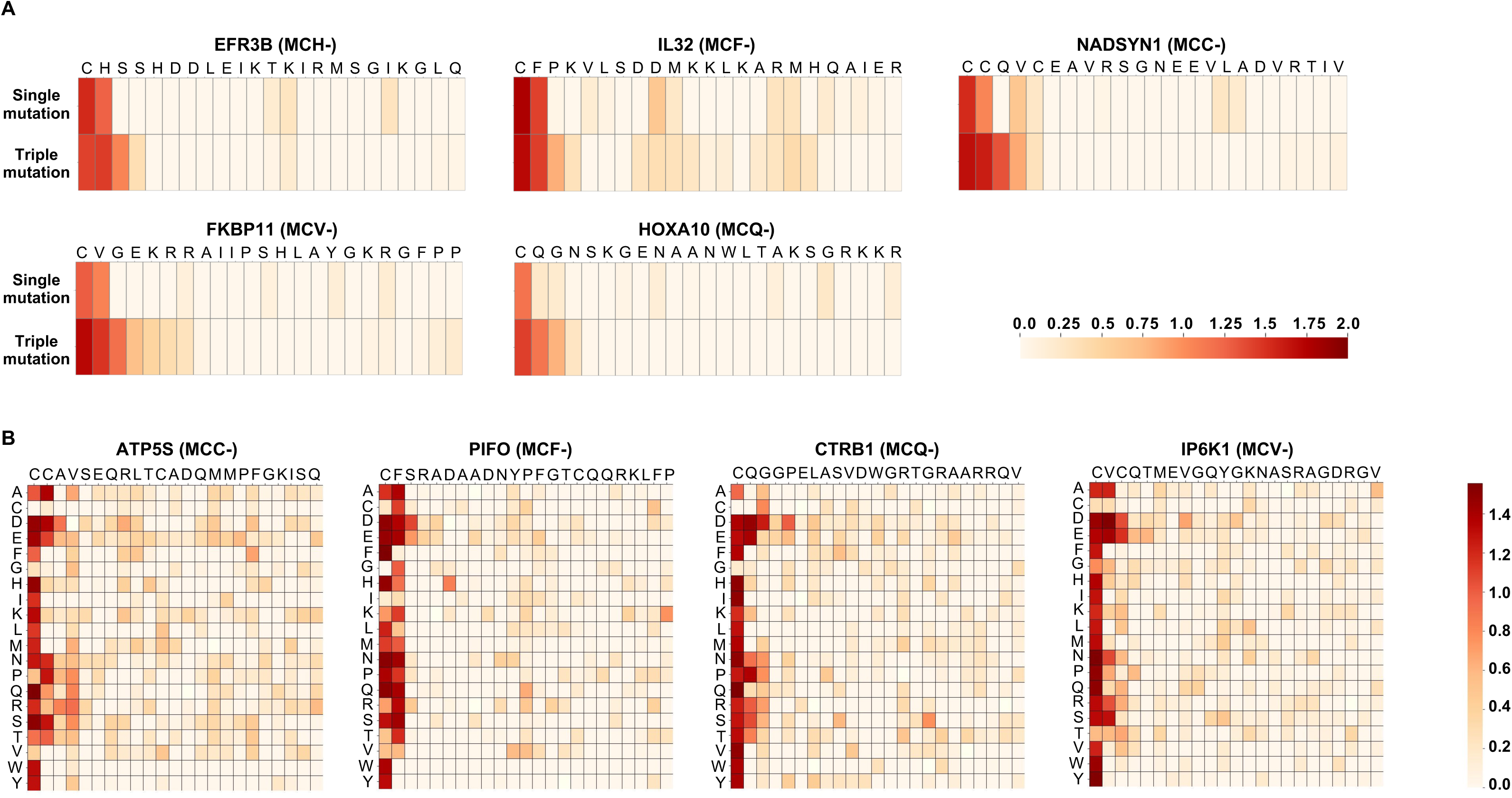
Specificity of the Cys-Arg/N-degron pathway revealed through mutagenesis screening (A) Scanning mutagenesis experiment of various N-terminal cysteine commencing peptides. For each of the indicated genes, data is presented for mutagenesis of single residues (top) or three consecutive residues (bottom). In each case, darker colors represent a greater degree of stabilization conferred by the mutation. (B) Saturation mutagenesis of four representative substrate examples with N-terminal cysteine. In each case, darker colors represent a greater degree of stabilization conferred by the mutation or N-terminal single amino acid addition. Gene’s name is indicated at the top and a universal scale of stabilization is shown on the right.

To gain further insight into the specific amino acids and positions contributing to peptide stability, we conducted saturation mutagenesis experiments, replacing each amino acid with the remaining 19 amino acids (Supplemental Table S2). Our results elucidated two key observations: first, Cys at the second position is most critical for substrate instability and cannot be substituted by any other residues (besides Gly in some cases, as Gly also serves as a potent N-degron (12)); second, the presence of residues such His, Phe, Cys, Val, Ile and Trp at the third position significantly contributed to peptide instability and can often be substituted for one another without disrupting stability (Figure 2B). These results are in line with enrichment of these residues in ATE1 peptide substrates (Figure 1B).

Overall, our findings underscore the significance of N-terminal Cys and neighboring residues, particularly at the penultimate position, in regulating peptide instability and susceptibility to degradation pathways.

### Oxygen- and ADO-dependent vs. ADO-independent mechanisms of stability regulation

Previously, it was demonstrated that following iMet cleavage, N-terminal cysteine residue can undergo N-terminal arginylation following chemical or enzymatic oxidation by ADO (2,14,17). To confirm whether the identified MC-substrates are regulated by ADO, peptide GFP-fusions with varying amino acid compositions at the third position were tested in ADO KO cells (Supplemental Figure 1B). The results revealed that all ATE1-positive substrates (“ATE1 positives”), including the positive controls RGS5 and IL32, were stabilized in ADO-deficient cells, with the exception of NADSYN1 (MCC-, Figure 3A). In contrast, MC-commencing peptides that were not identified as ATE1 substrates in the screen (“ATE1 negatives”) showed no stability change in response to ablation of ADO. In line with previous observations for established substrates like RGS5, the newly discovered peptide examples also exhibited substantial stabilization under hypoxic conditions as revealed by western blots (Figure 3B). Notably, in these assays HIF1α serves as another positive control substrate, stabilized under oxygen-deprived conditions (Figure 3B).

**Figure 3.**
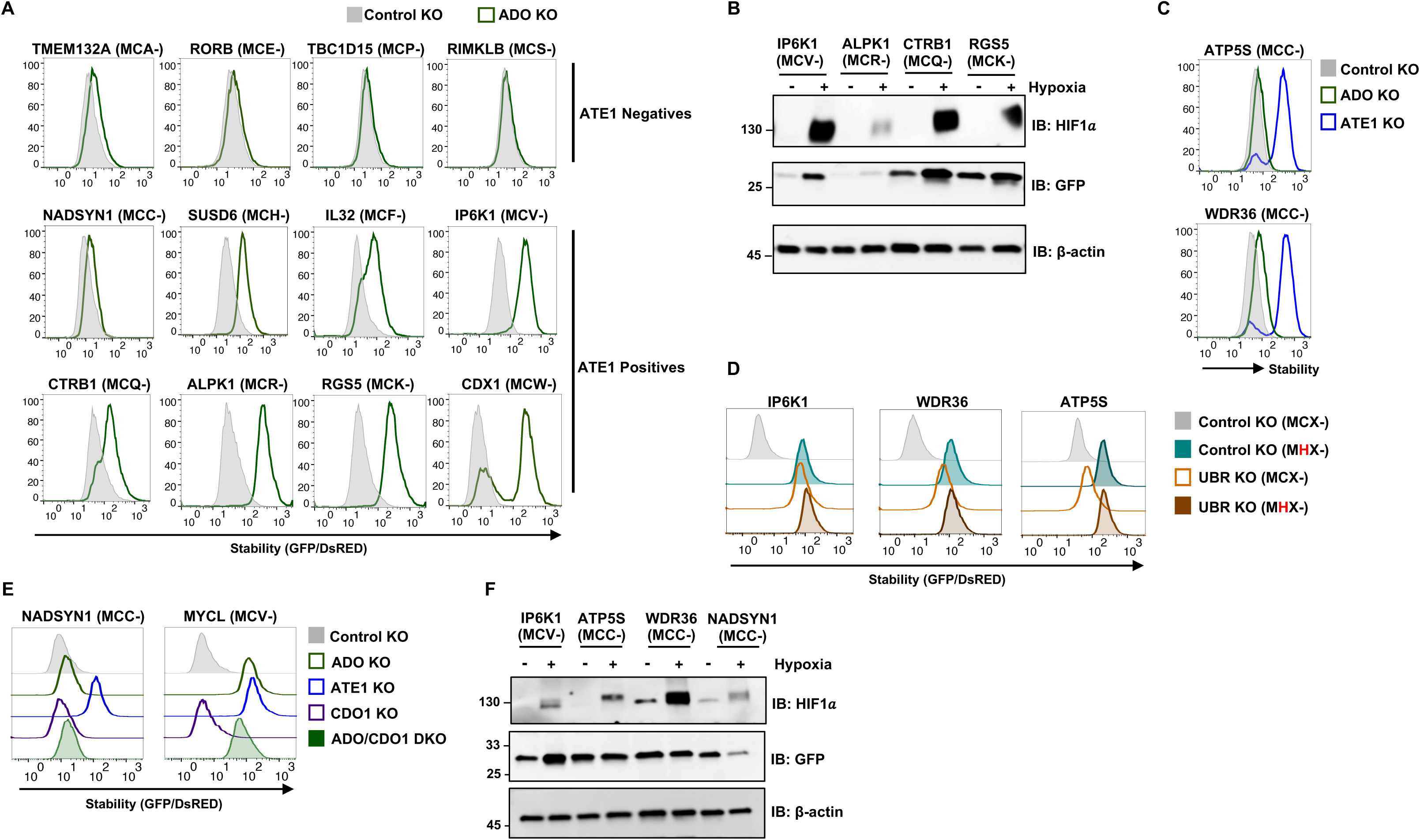
Analysis of oxygen-dependent and -independent regulation of N-terminal Cysteine substrates (A) Stability analysis of the indicated ATE1 negative and positive N24mer–GFP fusions assessed in control and ADO KO cells by flow cytometry. (B) HEK293T cells expressing N24mer–GFP fusions were incubated in hypoxic chamber for 16 h. Protein samples were then collected and analyzed by immunoblot with antibodies to GFP to detect peptide-GFP and β-actin was used as a loading control. Stabilization of endogenous HIF1α served as a positive marker for hypoxia stress response activation. RGS5 N24mer-GFP was used as positive control for Cys-commencing peptide-GFP fusion. (C) Stability analysis of N24mer–GFP fusions starting with MCC-motif in control, ADO and ATE1 KO cells by flow cytometry. (D) Stability analysis of WT (MCX-) or mutant (MHX-) peptide GFP-fusions expressed in control or UBR KO cells. (E) The stability of the indicated peptide GFP-fusions was analyzed in control, ADO, ATE1, CDO1 and ADO/CDO1 double KO cells. (F) Stability analysis in response to hypoxia of the indicated peptide GFP-fusions analyzed by immunoblot with the indicated antibodies.

We were specifically intrigued by the fact that NADSYN1 (MCC-) did not stabilize in ADO1 KO cells. Upon testing additional GFP-fusion peptides containing the MCC-motif (ATP5S, WDR36), it became clear that although all are substrates of the Arg/N-degron pathway, with their stability regulated by ATE1 and UBR, they are not subjected to regulation by ADO (Figure 3C). Notably, the the two adjacent Cys residues are the only ones shown to play a critical role in peptide turnover (Figure 2, see NADSYN1, ATP5S). To confirm that the degradation of these peptides depends on the N-terminal cysteine, we mutated Cys2 to histidine. While the wild type (WT) peptide was stabilized upon UBR ablation, the mutant peptide displayed substantial increased stability in control KO cells with no further stabilization observed in UBR KO cells (Figure 3D). This data underscores the critical role of the N-terminal MCC-degron for these substrate instability. To explore the possibility of another thiol dioxygenase catalyzing the oxidation of N-terminal Cys in these substrates, we generated KO cells of the only other known mammalian thiol dioxygenase, CDO1 (Supplemental Figure 1C). However, similarly to ADO KO, CDO1 deficiency as well as double KO of both dioxygenases ADO and CDO1 failed to induce any noticeable impact on stability of NADSYN1 (MCC-) (Figure 3E). Altogether the data suggest that MCC-commencing peptides are not subjected to regulation by ADO or CDO1. It is noteworthy that N-terminal cysteine can undergo spontaneous oxidation to sulfinic or sulfonic acid forms in the presence of potent oxidants like hydrogen peroxide and nitric oxide (NO). While this non-enzymatic modification could theoretically be inhibited in anaerobic conditions, intriguingly, hypoxia had no noticeable effect on MCC-peptide stability, while it stabilized IP6K1, an MCV-containing peptide (Figure 3F). This finding suggests that while ADO predominantly catalyzes cysteine oxidation to allow regulation of MC-substrates by ATE1 and Arg/N-degron pathway, alternative mechanisms may facilitate ATE1 recognition of unoxidized cysteine residues in MCC-commencing substrates. These pathways highlight additional complexity in Cys-Arg/N-degron mediated regulation.

### IP6K1 and PIFO are novel substrates of oxygen-dependent N-degron pathway

Next, we aimed to identify novel full-length proteins regulated by oxygen-dependent mechanisms. The MC-peptidome encompasses all classes of proteins, including extracellular, membrane-bound, organelle-specific, and cytoplasmic proteins. Additionally, some MC peptides may represent unique isoforms that are not the predominant isoforms expressed in cells. Among the substrates identified in the ATE1 and UBR screens, we specifically focused on cytoplasmic proteins and those defined as the primary isoform in UniProt. From this list, we selected two proteins that met these criteria: Inositol Hexakisphosphate Kinase 1 (IP6K1) and Primary Cilia Formation Protein (PIFO). Full-length versions of these proteins were cloned as GFP fusions, and their stability was analyzed in ADO, ATE1, and UBR KO cells, with IL32 serving as a positive control full-length protein. Flow cytometry revealed that both IP6K1 and PIFO were stabilized in all backgrounds (Figure 4A) suggesting regulation of these proteins by Cys-Arg/N-degron pathway. To rule out potential artifacts from the GFP fusion, we used effective endogenous antibodies for IP6K1 and demonstrated increased protein levels detected by western blotting (Figure 4B). For PIFO, which lacks available antibodies, we cloned it as a C-terminal HA fusion and generated single-integrant expressing cells via lentiviral transduction. Similar to IP6K1, PIFO-HA also showed increased protein levels in all KO cells by immunoblotting (Figure 4B). To assess oxygen-dependent regulation of these substrates, HEK293T and HeLa cells were exposed to hypoxia and the results demonstrated that both endogenous IP6K1 and PIFO-GFP proteins accumulated in cells exposed to hypoxia (Figure 4C) demonstrating hypoxia and Arg/N-degron regulation of these substrates.

**Figure 4.**
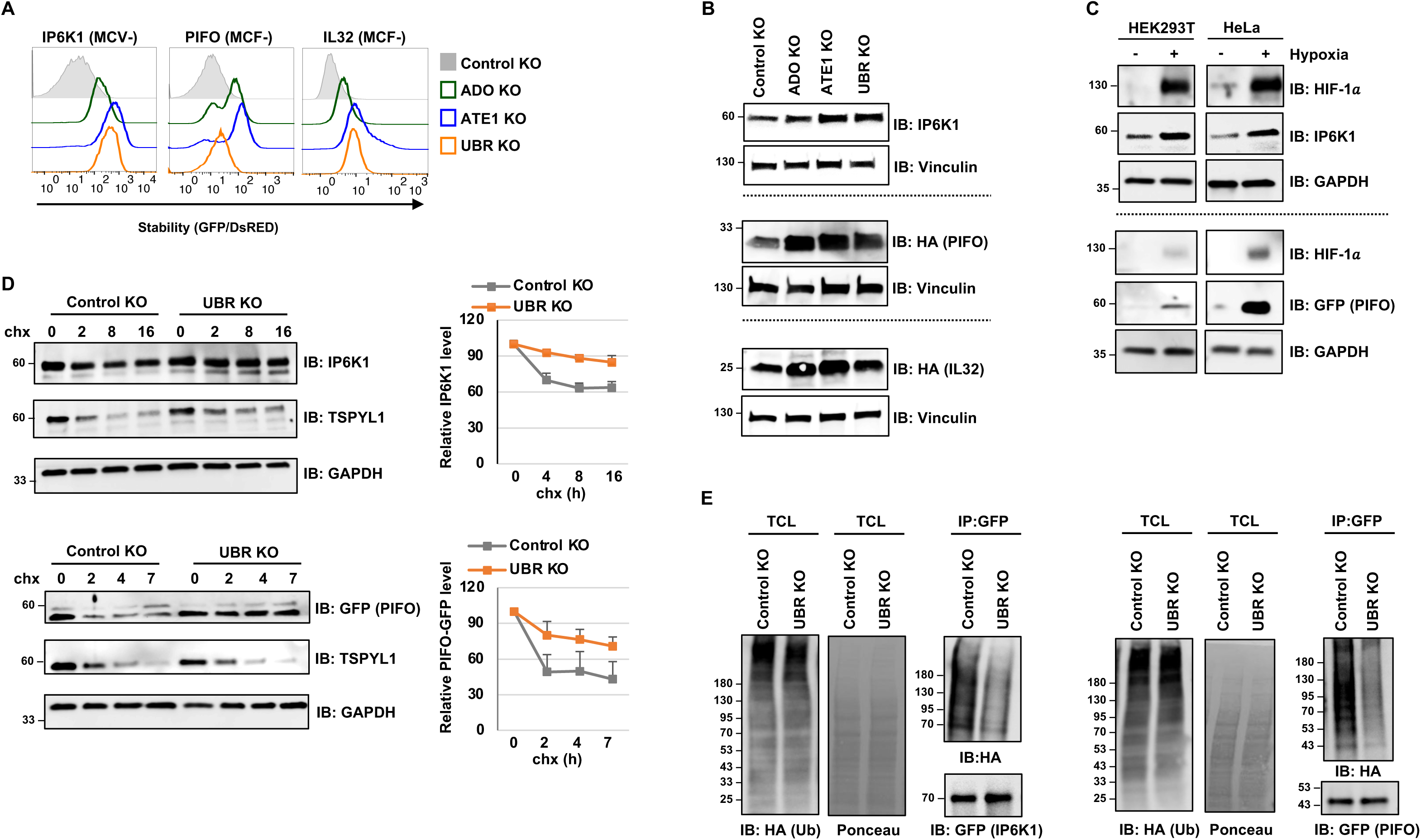
IP6K1 And PIFO are novel substrates of oxygen-dependent N-degron pathway (A) Stability analysis by flow cytometry of the indicated full-length GFP-fusion proteins in control, ADO, ATE1 and UBR KO cells. IL32-GFP was used as a positive control substrate. (B) Immunoblot of endogenous IP6K1, or PIFO-HA and IL32-HA stably expressed in HEK293T following 16 h exposure to hypoxia. Vinculin serves as a loading control. (C) Immunoblot of endogenous IP6K1 or PIFO-GFP stably expressed in HeLa cells. The stabilization of HIF1α serves as a positive indicator of hypoxia stress response activation. GAPDH serves as a loading control. (D) Representative western blots of Cycloheximide (chx) chase assays done to monitor the turnover rate of endogenous IP6K1 and PIFO-GFP in control or UBR KO cells by immunoblot (left). Quantification of IP6K1 and PIFO-GFP chx assays normalized to β-actin (right). TSPYL1 was used as a positive control for a short-lived protein during chx treatment. (E) Immunoblot of HA-ubiquitin conjugates and full-length IP6K1 or PIFO GFP fusions in total cell lysate (TCL) or GFP immunoprecipitates (IP:GFP) using anti-HA or -GFP antibodies, respectively.

Using a Cycloheximide (chx) chase assay to assess protein turnover rates, we observed that IP6K1 undergoes slow degradation, with approximately 60% of its initial protein levels remaining after 16 hours of treatment; in UBR KO cells the degradation of IP6K1 was attenuated (Figure 4D). PIFO, tagged with GFP, exhibited a more rapid degradation profile, which was also reduced in UBR KO cells (Figure 4D). Lastly, *in vivo* ubiquitination state of IP6K1 and PIFO was investigated in WT and UBR KO cells, revealing notably reduced levels of ubiquitination for both substrates in UBR KO cells (Figure 4E). These findings confirm that IP6K1 and PIFO are novel substrates of Arg/N-degron pathway, regulated by oxygen-dependent modulation of their turnover rates.

### IP6K1 plays a significant role in the hypoxia stress response

The next step was to investigate the biological significance of N-terminal Cys containing proteins during hypoxia. PIFO is known to play a crucial role in ciliary assembly and function, as well as in signaling pathways related to the cell cycle and stress responses (36,37). According to the Human Protein Atlas (HPA) (38), PIFO expressed in cells involved in the formation and function of cilia, including embryonic node, endometrial ciliated cells, and spermatids. However, as previously reported (36), ectopic expression of PIFO in HEK293T or U2OS cells, both of which naturally lack PIFO expression, resulted in the activation of target genes *GLI1* and *PTCH1*, key components of the Hedgehog signaling pathway under normoxic condition (Supplemental Figure 2A, B). In HEK293T under hypoxic conditions, increased levels of PIFO due to inhibition of its degradation (Figure 4C) led to further increase in the accumulation of *GLI1* and *PTCH1* (Supplemental Figure 2B). The ATP5S-GFP protein, which is expressed at higher levels than PIFO as indicated by flow cytometry (Supplemental Figure 2C), was used as a negative control and did not induce *GLI1* or *PTCH1* expression under the same settings. These findings suggest that hypoxia-induced PIFO stabilization boosts Hedgehog signaling. The effects of this activation on cellular adaptation and metabolism during hypoxia are beyond the scope of this study and require further investigation to assess the functional impact of PIFO stabilization.

IP6K1 is a versatile enzyme that impacts various cellular processes through its role in generating inositol pyrophosphates (IPs) (27). IPs, such as IP7 and IP8, have one or more energy-rich pyrophosphate moieties that have a free energy of hydrolysis comparable to that of ATP. IP7 is also involved in several physiological functions, including insulin secretion and AKT signaling (39,40). Given the regulation of IP6K1 turnover under hypoxia and its role in cellular signaling, we explored its potential involvement in the hypoxic response.

Since IP6K1 protein levels increase under hypoxia, we examined the effects of its depletion. To this end, we generated two independent IP6K1 KO cell lines in HeLa cells (Supplemental Figure 1D,E) and assessed their proliferation under normoxic and hypoxic conditions using colony formation assay. While no significant difference in colony formation was observed under normoxia, IP6K1 KO cells showed a marked reduction in colony formation under hypoxia compared to control cells (Figure 5A,B). These findings suggest that IP6K1 deficiency markedly impairs cell proliferation and inhibits colony formation in hypoxic environments.

**Figure 5.**
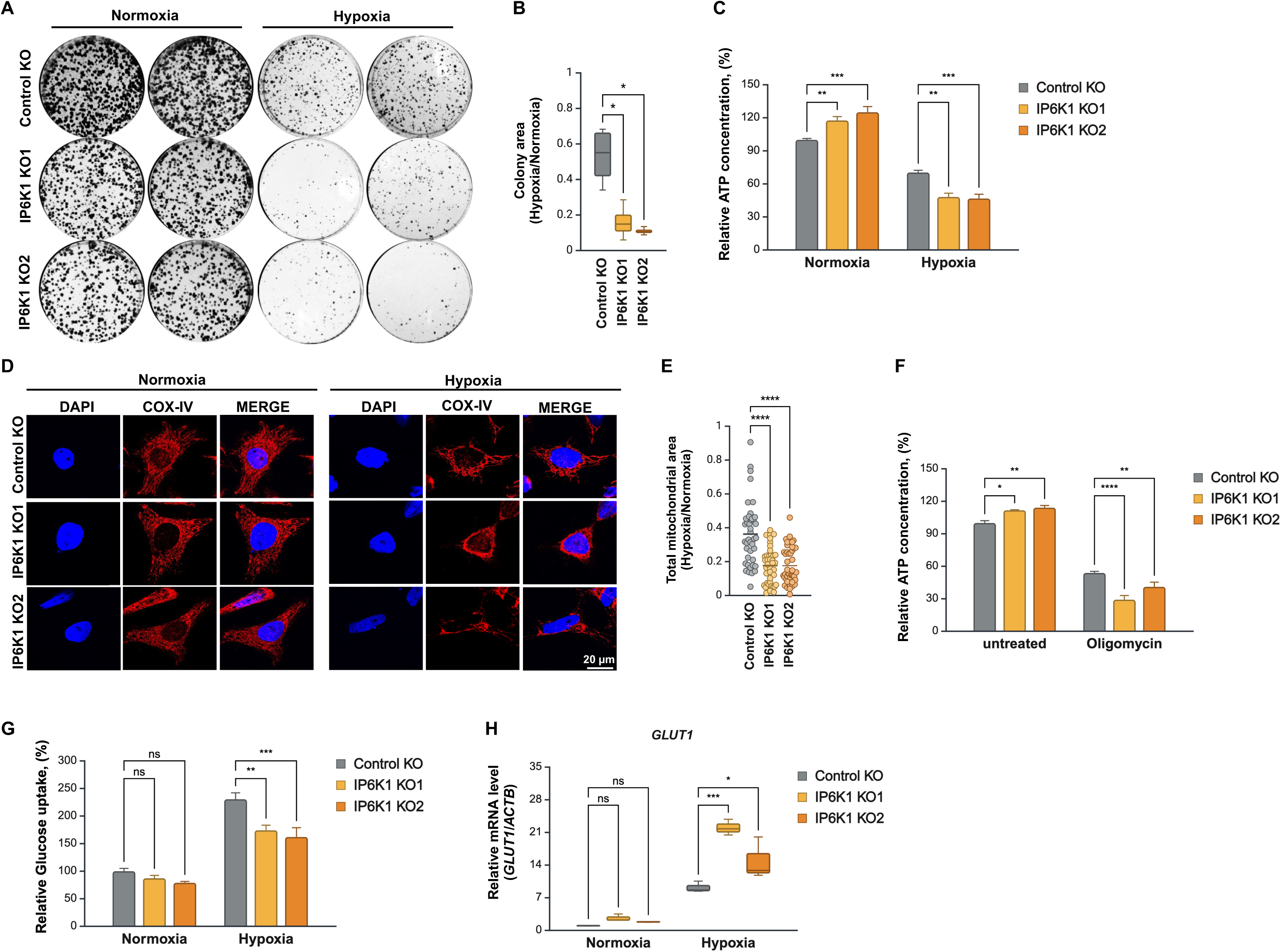
IP6K1 regulates metabolic reprogramming during hypoxia (A-B) Colony formation assay of two independent IP6K1 KO cell lines (sg1, sg2) compared to control KO HeLa cells grown in normoxic or hypoxic conditions for 12 days. (A) Representative images and (B) Quantitative analysis of colony formation assay from (A). (C) Intracellular ATP level measured in IP6K1 KO and control KO HeLa cells cultured in normoxic or hypoxic conditions for 16 h. (D-E) Mitochondria morphology was analyzed in IP6K1 and control KO HeLa cells under normoxic or hypoxic conditions. (D) Representative images of mitochondrial visualized by immunostaining with COX-IV antibody. (E) Quantitative analysis of mitochondrial morphology with ImageJ. DAPI (blue) marks the nucleus. Scale bar= 20 μM. (F) Intracellular ATP level in IP6K1 and control KO HeLa cells treated with 10 μM oligomycin for 16 h. (G-H) Glucose uptake (G) and mRNA level of *GLUT1* normalized to b-actin (H) in control or IP6K1 KO cells cultured in normoxic vs hypoxic conditions. Data presented as mean. Ns-no significance; *P < 0.05; **P < 0.01; ***P < 0.001; ****P < 0.0001.

IP6K1 plays a crucial role in maintaining cellular metabolic balance and ATP production (41), presumably by regulating phosphate export through the inhibition of XPR1, a phosphate exporter, which leads to increased intracellular ATP levels (42). Consistent with these findings, IP6K1 KO cells exhibited elevated ATP levels under normoxic conditions (Figure 5C). However, during hypoxia, ATP levels decreased by approximately 30%, in control cells, whereas IP6K1 KO cells experienced a more pronounced reduction of 50% (Figure 5C).

Given the link between hypoxia and mitochondria functions (43,44), we speculated that the effect IP6K1 on energy homeostasis might be elicited by mitochondrial dysfunction and disruption of oxidative phosphorylation. To explore this further, we examined mitochondria morphology by confocal microscopy. Hypoxia was found to reduce the overall mitochondrial area, with mitochondria accumulating in the perinuclear region (Figure 5D, E). In the absence of IP6K1, there was a marked decrease in mitochondrial area, accompanied by an even greater perinuclear clustering (Figure 5D, E), indicating that the loss of IP6K1 contributes to mitochondrial dysfunction.

Building on these morphological changes, we next evaluated the impact of mitochondrial dysfunction on cell survival by culturing cells in galactose, a substrate that can only be metabolized by respiring mitochondria, and analyzed colony formation over 12 days. IP6K1 KO cells formed fewer colonies under these conditions, suggesting that cells lacking IP6K1 have compromised mitochondrial functionality (Supplemental Figure 3A).

This mitochondrial dysfunction prompted us to further investigate how metabolic adaptation might be affected in the absence of IP6K1. Hypoxia was shown to induce a metabolic shift from mitochondrial respiration to anaerobic glycolysis (45). To assess glycolytic ATP production in the absence of IP6K1, cells were treated with oligomycin, an inhibitor of ATP synthesis via oxidative phosphorylation (OXPHOS). The results revealed that IP6K1 KO cells exhibited lower ATP levels in response to oligomycin treatment (Figure 5F), mirroring the decrease seen during hypoxia conditions (Figure 5C), suggesting that IP6K1 plays a role in glycolysis during hypoxia. Additionally, glucose uptake, which is typically enhanced by hypoxia, was approximately 30% lower in IP6K1 KO cells compared to control cells (Figure 5G), indicating that IP6K1 depletion restricts glucose consumption under hypoxic conditions. Since hypoxia stimulates glucose transport through increased transcription of the *GLUT1* glucose transporter gene (46), we evaluated *GLUT1* expression and found it was more strongly induced in IP6K1 KO cells (Figure 5H), likely reflecting a compensatory feedback mechanism to counteract reduced glucose uptake in these cells (Figure 5G). The altered mitochondria function was also evident from the expression patterns of OXPHOS and glycolytic genes, which were upregulated in IP6K1 KO (Supplemental Figure 3B,C) likely due to impaired glucose uptake under hypoxia.

Taken together these findings suggest that IP6K1 is critical for maintaining mitochondrial function and facilitating metabolic adaptation during hypoxia. Thus, increasing IP6K1 protein levels during hypoxia by preventing its degradation via the Cys-Arg/N-degron pathway could enhance its function under these conditions.

## Discussion

Hypoxia activates signaling pathways that enable cell survival and adaptation under stress (47,48). A recent study by Tian et al (49), compared two oxygen-sensing regulatory systems and suggested that enzymatic protein oxidation by ADO coupled with proteolysis control of HIF1α may serve as first-line responses to hypoxia. While the induction of HIF1α protein is rapid during hypoxia due to inhibited proteolysis, the accumulation of HIF1α target genes is delayed as transcriptional adaptions take longer than posttranslational events. In contrast, direct regulation of N-terminal cysteine-containing proteins through the Cys-Arg/N-degron pathway can provide a more immediate response to hypoxic conditions. Unlike HIF1α-dependent regulation, protein substrates of the N-degron pathway are not regulated by feedback inhibition, enabling them to remain stable and functional during prolonged hypoxic stress (49). Here we systematically searched for proteins regulated at their turnover rate in response to hypoxia as part of the Cys-Arg/N-degron pathway.

Proteins starting with an MC-sequence are underrepresented in the both plant and human genome (12,50), likely due to the robustness of Cys-Arg/N-degron pathway toward these substrates and evolutionary pressure against these sequences, as also seen for other terminal degrons (12,25). Our scanning mutagenesis experiment demonstrated that substrate selectivity in Cys-Arg/N-degron pathway extends beyond N-terminal Cys and that there is a pronounced preference for hydrophobic and positively charged residues immediately following the N-terminal Cys. These ATE1-dependent motifs resemble, although not fully match, the motifs recognized by ADO *in vitro* (24); An *in vitro* peptide activity assay showed that ADO exhibited the highest activity towards peptides with basic (Lys/Arg/His) or aromatic (Phe/Trp) residue following the Cys (24). The ability of ADO to interact with amino acids of varying sizes and properties can be explained by structural analyses of human ADO that revealed an open catalytic cavity and flexible loops at the active site entrance (51). These structural features allow ADO to accommodate a diverse range of substrates enabling it to function effectively as an oxygen sensor. The discrepancy between our findings and those of Heathcote et al. may be due to the *in vitro* nature of their assays, which do not fully represent *in vivo* conditions. Additionally, the lower throughput of *in vitro* assays limits the coverage of substrates tested, potentially overlooking some consensus motifs. Additionally, while ADO specificity was revealed through *in vitro* assays, our GPS assays specifically identify MC-substrates regulated by stability. Protein stability via the Cys-Arg/N-degron pathway is regulated not only by ADO but also by ATE1, and UBR proteins. Thus, the substrate specificity of ADO may not perfectly align with the specificity required by the downstream regulators ATE1 and UBRs leading to a different substrate range than what ADO activity alone would suggest. Moreover, it has been shown that ADO and the N-terminal acetyltransferase NatA have distinct preferences for N-terminal Cys substrates protecting against opposing modifications *in vitro* (24). Therefore, regulation of protein stability is complex and involve a balance of stabilizing or destabilizing post-translational modifications that play a critical role in determining protein stability.

ATE1 predominantly catalyzes the addition of Arg to N-terminally exposed acidic residues such as aspartate (Asp), glutamate (Glu), and oxidized cysteine (Cys, sulfinic/sulfonic acid) (2,17,52). However, significant gap remains in understanding ATE1 substrate preferences. Our results demonstrate that ATE1 can regulate the stability of specific class of N-terminal Cys proteins, those that encode MCC-motif independently of ADO. Although N-terminal cysteine can be oxidized directly by nitric oxide (NO) (53) evidence further suggests that the MCC-degron is not regulated by hypoxia. Does the presence of two vicinal cysteine residues can increase the likelihood of oxidation by nitric oxide (NO) or whether ATE1 might recognize unmodified cysteine residues remains an open question.

Our study extends beyond revealing the substrate specificty of Cys-Arg/N-degron pathway to explore novel substrates and their biological implications. We identified IP6K1 as a key metabolic enzyme that requires tight regulation of its degradation to maintain cellular homeostasis. Previously, it was shown that elevated IP6K1 activity, linked to increased IP7 production, disrupts DNA repair and protein phosphorylation (29,54). Additionally, IP6K1 impairs insulin signaling (28), which may contribute to insulin resistance and metabolic disorders such as obesity and diabetes (55). In contrast, IP6K1 deletion or inhibition reduces cell invasiveness and migration, offering protection against carcinogenesis (26). This altogether highlights the need for its tight regulation and positionin IP6K1 as a potential therapeutic target. Given the role of IPs in regulating metabolic balance between glycolysis and oxidative phosphorylation (41,56), we investigated the role of IP6K1 in metabolic control under low-oxygen conditions. Depletion of IP6K1 resulted in reduced glucose uptake in hypoxia, impairing glycolytic ATP production consistent with the fact that cells rely heavily on glycolysis for ATP production when oxidative phosphorylation is restricted during hypoxia. This also limited the availability of key metabolic intermediates necessary for nucleotide, lipid, and amino acid biosynthesis, thereby impairing cell proliferation during hypoxia (45,47,48). To cope with metabolic stress, hypoxia induces the transcription of glycolytic enzymes and glucose transporters. Although IP6K1 KO cells show increased transcript levels of GLUT1, OXPHOS, and glycolytic genes, their failure to uptake glucose—normally enhanced during hypoxia— disrupts this adaptive mechanism. Hypoxia-mediated adaptations involve changes in the regulatory control of key molecular components of metabolic pathways centered on the mitochondria, as well as dynamic changes in the morphology, mass and subcellular localization of mitochondria themselves (57,58). Consistent with this, we observed that IP6K1-deficient cells exhibited altered mitochondrial morphology. Moreover, these cells showed reduced survival, reinforcing the importance of IP6K1 in cellular homeostasis.

Oxygenation levels vary across tissues due to variations in metabolic demand, vascularization, and environmental conditions. While “normoxia” refers to atmospheric oxygen (about 20%), “physioxia” in the body ranges from 11% to 1% depending on the tissue (59). This variability underscores the importance of considering tissue-specific oxygen levels in cellular studies. Future research comparing the turnover of MC-starting proteins across tissues could reveal their roles in oxygen homeostasis and uncover tissue-specific regulatory mechanisms, enhancing our understanding of cellular adaptation to oxygen fluctuations in health and disease.

## Materials and Methods

### Cell culture

HEK293T (ATCC CRL-3216), HeLa cells (a gift from Kimchi A., Weizmann Institute of Science) and U2OS cells (A gift from Shav-Tal Y. lab, Bar-Ilan university), were grown in Dulbecco’s Modified Eagle’s Medium (DMEM) (Life Technologies) supplemented with 10% fetal bovine serum (FBS) (Gibco) and penicillin/streptomycin (Life Technologies). Cells were maintained at 37 °C under an atmosphere of 5% CO2 in air. Hypoxia was induced by incubations within an Hypoxia Incubator Chamber (# 27310; STEMCELL). The proteasome inhibitor Bortezomib (#A2614; APExBio) was used at a final concentration of 1 μM and Cycloheximide (#A8244; APExBio) was used at a final concentration of 100 μg/ml.

### Transfection and lentivirus production

Lentivirus was generated through the transfection of HEK293T cells using PolyJet In Vitro DNA Transfection Reagent (#SL100688; SignaGen Laboratories). Cells seeded at ∼80% confluency were transfected as recommended by the manufacturer with the lentiviral transfer vector plus four plasmids encoding Gag-Pol, Rev, Tat, and VSV-G. The lentiviral supernatants were collected 48 h later. Transduction of target cells was achieved by adding the virus in the presence of 8 μg/ml hexadimethrine bromide (Polybrene) (# H9268; Sigma-Aldrich).

### Plasmids

Open reading frames (ORFs) encoding IP6K1, PIFO, IL32 and ATP5S were obtained from the Ultimate ORF Clone collection (Thermo Fisher Scientific). ORFs were amplified by PCR to include *XhoI* and *BstBI* sites and were cloned using the Gibson assembly method (NEBuilder HiFi Cloning Kit) into pHAGE-GFP-IRES-DsRed vector N-terminal to GFP. To monitor ORFs levels by western blot, ORFs were amplified by PCR to include an HA epitope at their C-terminus and subcloned into the lentiviral pHAGE vector that also contains IRES-DsRed cassette to monitor equal expression of the constructs in various cell lines by flow cytometry.

For individual CRISPR/Cas9-mediated gene disruption experiments, the lentiCRISPR v2 vector was used (#52961; Addgene). Oligonucleotides encoding the top and bottom strands of the sgRNAs were synthesized (IDT), annealed, and cloned into the lentiCRISPR v2 vector as previously described (60).

Nucleotide sequences of the sgRNAs used were:

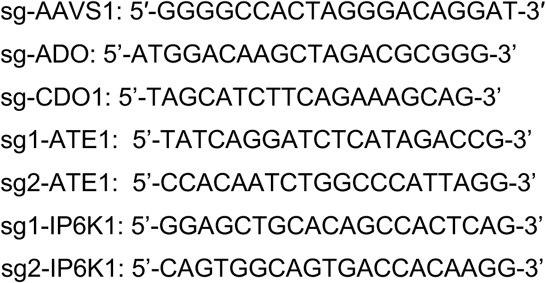

### Generation of CRISPR/Cas9 knockout cells

Lentivirus was generated through the transfection of HEK293T with lentiCRISPR v2 as explained before. Briefly, 48 h following transduction cells were selected with puromycin to eliminate non transduced cells. Seven days post transduction, genomic DNA of transduced cells was extracted, PCRs were performed to amplify ∼500 flanking the edited site followed by Sanger sequencing. Inference of CRISPR Edits (ICE) CRISPR Analysis Tool was used to analyze efficiency of editing (Synthego Performance Analysis, ICE Analysis. 2019. v3.0.). UBR1/2/4 KO clone #2 (12) is used in this study as it gave the strongest stabilization effect of known UBR substrates

### Global protein stability (GPS) screen

The generation of a GPS lentiviral vector expressing a N-terminal peptide library was described previously (12). Briefly, the oligonucleotide pool synthesized by Agilent Technologies, was cloned into Ub-GPS vector using Gibson assembly downstream to ubiquitin and followed by GFP. For the scanning mutagenesis screen, oligonucleotides were synthesized to introduce mutations at individual residues or sets of three consecutive residues. Amino acids were replaced with ones possessing different characteristics from the original residues to disrupt potential degrons (e.g., large non-polar to small polar, acidic to basic) (31). For saturation mutagenesis, each amino acid across all positions in selected peptides was mutated to all other 19 amino acids. The library was packaged into lentiviral particles and introduced into HEK293T cells at a multiplicity of infection of ∼0.2 (achieving approximately 20% DsRed+ cells). After selection with Hygromycin (100 μg/ml) cells were partitioned into four bins of equal size based on the stability of the GFP fusion (GFP/DsRed ratio). Genomic DNA was extracted from cells collected from each of the bins using Gentra Puregene Cell Kit (#158767; Qiagen) and the fusion peptides were amplified by PCR (Q5 Hot Start Polymerase; NEB) using primers binding in constant regions flanking the N24mer peptide for the first PCR reaction, followed by a second PCR reaction to add Illumina indexes and P5 and P7 adaptors. Samples to be multiplexed were then pooled, purified on an agarose gel (QIAEXII Gel Extraction Kit, #20051; Qiagen), and sequenced on an Illumina NextSeq instrument.

*Data analysis:* was performed as described previously (12,31). Raw Illumina reads derived from each GPS bin were first trimmed of constant sequences derived from the Ub-GPS vector backbone using Cutadapt (61) and count tables were generated from reads that aligned perfectly to the reference sequence. Following correction for sequencing depth, and based on the read counts of each peptide in every bin, the protein stability index (PSI) metric was calculated for each peptide-GFP fusion. The PSI score is given by the sum of multiplying the proportion of reads in each bin by the bin number thus yielding a stability score between 1 (maximally unstable) and 4 (maximally stable):

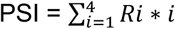

(where *i* represents the number of the bin and *R_i_* represents the proportion of Illumina reads present for a peptide in that given bin *i*).

Read counts and associated stability score for each peptide-GFP fusion in the ATE1 KO screen are detailed in Supplemental Table 1. Candidate ATE1 substrates were defined as if they showed ΔPSI≥0.3 (PSI_ATE1_-PSI_Control_) for both replicates.

For classification of UBR substrates from the Ub-GPS screen detailed by Timms et al (12): a ΔPSI score was generated for each peptide-GFP fusion reflecting the difference in raw PSI scores between control KO sample and UBR KO clones #1-3. Peptide-GFP fusions were defined as UBRs substrates if they were stabilized ≥0.3 PSI units in at least one UBR KO clone. In addition, GPS screen profiles for each of representative Ub-GPS N24mers commencing with Cys showing the distribution of Illumina sequencing reads across the bins in control cells versus three independent UBR1/2/4 KO cells were plotted. Those with a positive median geometric mean stability of the peptide in UBR KO cells compared to WT cells (indicating a stabilization shift in UBR KO cells) were filtered as substrates.

For the mutagenesis screens-a ΔPSI score was generated for each peptide-GFP fusion reflecting the difference in raw PSI scores between wild type peptides and mutants. These values are detailed in Supplemental Table 2 and presented as heatmaps in Figure 2.

### Flow cytometry

Analysis of HEK293T cells by flow cytometry was performed on CytoFlex (Beckman Coulter) instrument and the resulting data were analyzed using FlowJo software. Cell sorting was performed on BD FACS Aria II (Becton Dickinson).

### Immunoblotting

Protein samples were collected in lysis buffer (10 mM NaPO4, 100 mM NaCl, 5 mM EDTA pH 8, 1% Triton X-100, 0.5% Deoxycholic acid sodium salt, 0.1% SDS) supplemented with Halt Protease and Phosphatase Inhibitor Cocktail (Thermo Fisher Scientific) and lysed for 30 min in +4^0^C. Protein concentration was determined by a standard Bradford assay (#500-0006; Bio-Rad), a linear bovine serum albumin (BSA) calibration curve, and an Epoch microplate spectrophotometer. 30 μg of total cell extract media were subsequently resolved by SDS-PAGE (Mini-PROTEAN TGX Precast Protein Gels; Bio-Rad) and transferred to a nitrocellulose membrane (Trans-Blot Turbo System; Bio-Rad) which was then blocked in 10% nonfat dry milk in PBS-T. The membrane was incubated with primary antibody overnight at 4°C, and then, following three washes with PBS-T, HRP-conjugated secondary antibody was added for 1 h at room temperature. Following a further three washes in PBS-T, reactive bands were visualized using SuperSignal West Femto chemiluminescence substrate (#34095; Pierce) or an EZ-ECL (#20-500-171; Biological Industries) for 5 min using the ImageQuant TL software v8.2 on Amersham Imager 680 (Cytiva). Primary antibodies used: HIF-1α (# A300-286A, FORTISLIFE - Bethyl), GFP (Abcam, ab290), IP6K1 (A305-628A-T; FORTISLIFE - Bethyl) TSPYL1 (#ab95943; Abcam), HA-Tag (C29F4) (#3724; Cell Signaling Technology), Vinculin (#V9131; Sigma-Aldrich), β-actin (#4967; Cell Signaling Technology), GAPDH (14C10) (#2118; Cell Signaling Technology). HRP-conjugated goat anti-rabbit and anti-mouse IgG secondary antibodies were obtained from Jackson ImmunoResearch (#111-035-003, #115-035-003, respectively). Densitometric analysis was performed using ImageJ software (NIH) and values were presented relative to housekeeping protein.

### Cycloheximide chase assay

Following treatment with 100 μg/ml Cycloheximide, cells were harvested at the indicated time points and subjected to Western blot as explained before.

### Analysis of ubiquitination

HEK293T cells stably expressing ORF fused to GFP were grown in 10 cm plates and transfected with HA-ubiquitin plasmid (25). 48 h post transfection, cells were treated with bortezomib (1 μM, 5 h), and then were lysed in ice-cold lysis buffer (10 mM NaPO4, 100mM NaCl, 5mM EDTA pH 8, 1% Triton X-100, 0.5% Deoxycholic acid sodium salt, 0.1% SDS) supplemented with HaltTM Protease and Phosphatase Inhibitor Cocktail (Thermo Scientific) and 50 µM of the deubiquitinating enzyme inhibitor PR-619 (#662141; EMD Millipore) for 30 min on ice. Protein concentration was determined by Bradford assay and equal amounts were taken for immunoprecipitation experiment by incubation for 2 h with 20 μl beads coated with anti-GFP (GFP-Trap_MA magnetic agarose beads (ChromoTek GmbH)). The beads were then washed three times with stringent washes in denatured conditions using wash buffer (10 mM NaPO4, 300 mM NaCl, 5mM EDTA pH 8, 1% Triton X-100, 0.5% Deoxycholic acid sodium salt, 0.1% SDS) supplemented with HaltTM Protease and Phosphatase Inhibitor Cocktail and 50 µM PR-619 before bound proteins were eluted upon incubation with SDS-PAGE sample buffer (95°C, 10 min). SDS-PAGE and immunoblot was done as explained before.

### Immunocytochemistry

Cells were fixed with 4% paraformaldehyde and then permeabilized with 0.2% Triton X-100. After blocking with 3% bovine serum albumin (BSA) in PBS, mitochondria were stained with COX-IV antibody (#4850; Cell Signaling Technology). Following three washes with PBS + 0.1% Tween-20 (PBS-T), DAPI staining (#D9542; Sigma-Aldrich) was added for 1 min. Finally, coverslips were mounted onto slides using mounting media (#F6182; Sigma-Aldrich) prior to imaging. Cells were imaged with Leica Stellaris 5 confocal microscope using LASX software and a 63× oil lens /1.4 N.A. UPlanSApo objective (Olympus). Analysis was done by in ImageJ using the Mitochondria Analyzer plugin (62).

### ATP and Glucose uptake measurement assays

ATP level was quantified using CellTiter-Glo assay (#G7571; Promega) according to manufacture protocol. Briefly, for ATP detection, for each well containing 500 μl media, 500 μl of CellTiter-Glo reagent was added. The luminescence levels from 200 μl per well were recorded after 10 minutes of incubation.

Glucose uptake was quantified using Glucose Uptake-Glo (#J1341; Promega) according to manufacture protocols. Briefly, for glucose uptake detection, cells were rinsed in warmed PBS and placed in 100 μL PBS in 24-well culture plate. 2-Deoxyglucose (1 mM) was added for 10 minute uptake. After stopping and neutralization, 2-deoxyglucose-6-phosphate detection reagent was added, supernatant was transferred to white 96-well plate (Costar) and luminescence was recorded at 30 minutes.

Normalization of cell number was done with Hoechst 33342 stain (62249, ThermoFisher Scientific) of 1 ng/ml final concentration. Fluorescence signal intensity was detected in plate reader at excitation ∼350 nm and emission ∼460 nm wavelength.

### Colony formation assay

To evaluate the ability of single cancer cells to form a colony, cells were plated at a concentration of 2,000 cells/well on six-well plates (day 0) in replicates and were grown in normoxic and hypoxic conditions. At day 10, cells were fixed with methanol:acetic acid (3:1) for 5 min. Colonies were stained with crystal violet working solution (0.1% in methanol) for 15 min. Colonies areas were measured for each well using ImageJ software (version 1.54 h; National Institutes of Health) and the ratio between hypoxia and normoxia was calculated.

### RT-qPCR

mRNA was extracted using RNEASY PLUS MINI KIT (74104; QIAGEN) and equal amounts were used for cDNA synthesis using the qScript cDNA Synthesis Kit (101414; Quantabio). qPCR analysis was performed using Applied Biosystems™ Power SYBR™ Green PCR Master Mix, on a CFX96 Touch Real-Time PCR Detection System (BioRad) using the ΔΔCt method. Levels of the housekeeping gene β-actin (*ACTB*) were used as a reference. Sequences for the primers used are as follows:

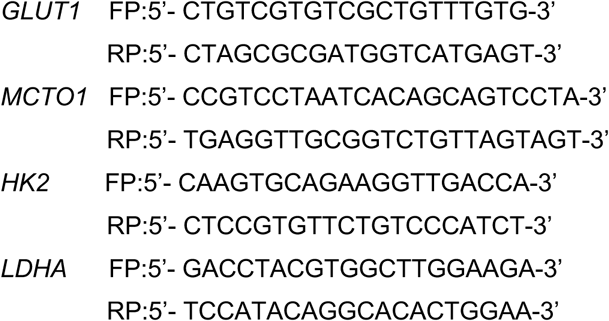

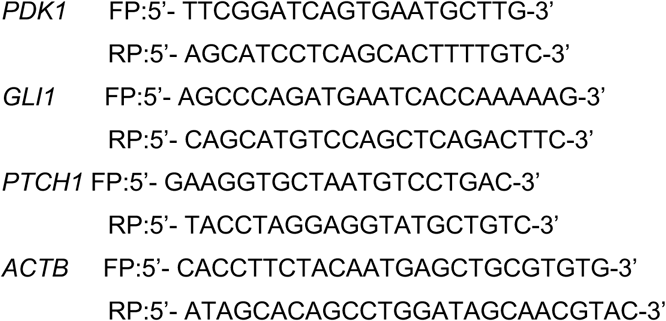

## Statistical analysis

To evaluate statistical significance of the enrichment of residues in position 3 in MC**X**-containing proteins a Z-score was calculated with the formula:

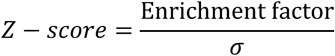

The Enrichment factor is the ratio of CX-occurrences to the total occurrences of X residue in position 3 in the proteome. σ is the standard deviation of CX-positive occurrences. Unless specified otherwise, data are expressed as means ± SEM.

Comparisons between groups were made via unpaired two-tailed Student’s t-test and two-way ANOVA. Statistical significance was set at P ≤ 0.05; ns indicates no significance, P > 0.05; *P < 0.05; **P < 0.01; ***P < 0.001; ****P < 0.0001.

## Acknowledgments

We thank Timms R.T. for assistance with the design and synthesis of the mutagenesis peptide library. This study was supported by by European Research Council (ERC-2020-STG 947709) and Israel Science Foundation (ISF Grants No. 2380/21 and 3096/21).

## Author Contributions

Conceptualization, A.B. and I.K.; Supervision, I.K.; Methodology, A.B. and I.K.; Software, A.B., S.B.-D. and I.K.; Formal analysis, A.B., S.B.-D. and I.K.; Investigation, A.B., Y.M. and I.K.; Writing-Original draft, A.B. and I.K.

## Competing Interest Statement

The authors declare that they have no conflict of interest.

## Supplemental Figure legends

**Supplemental Figure 1.**
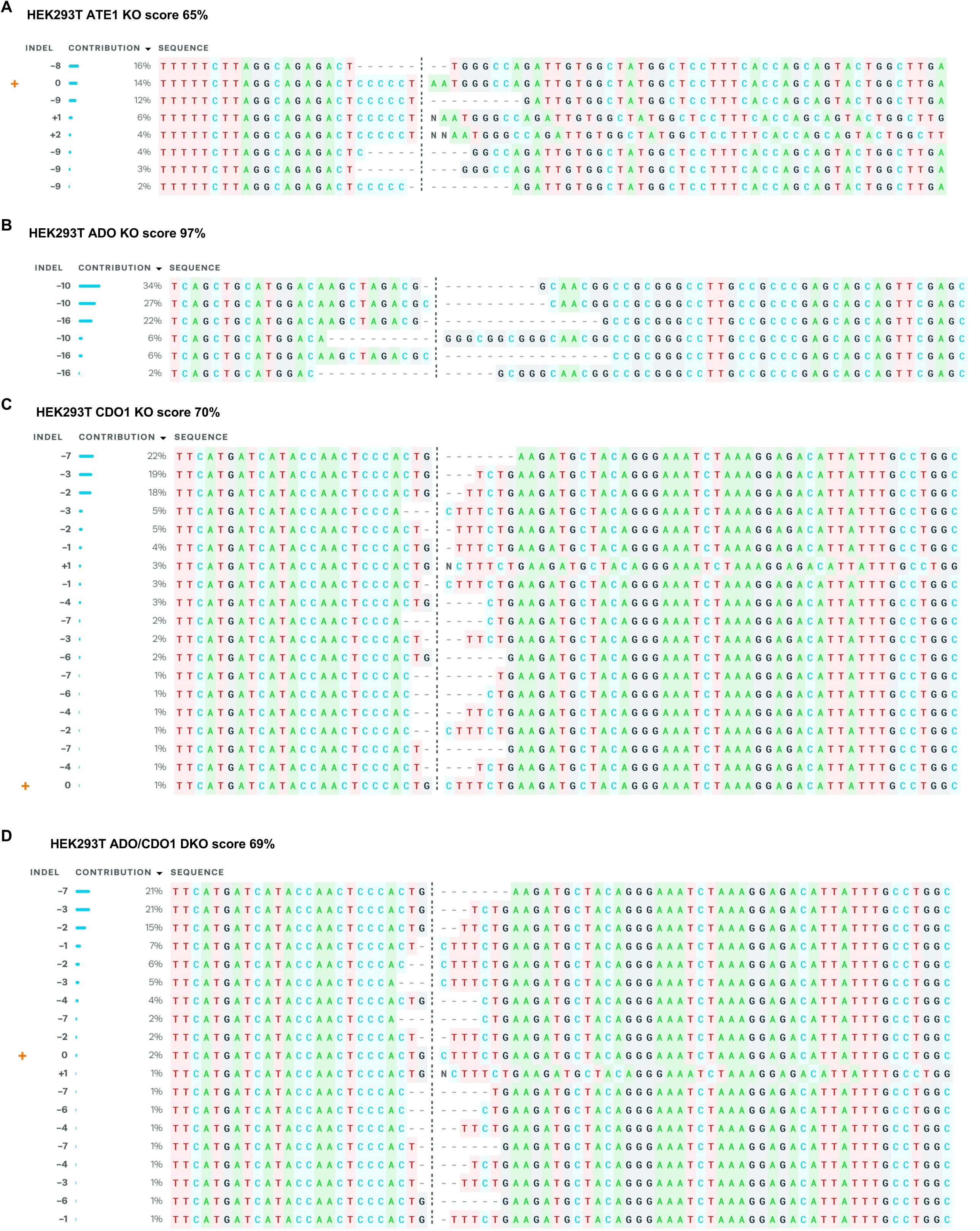
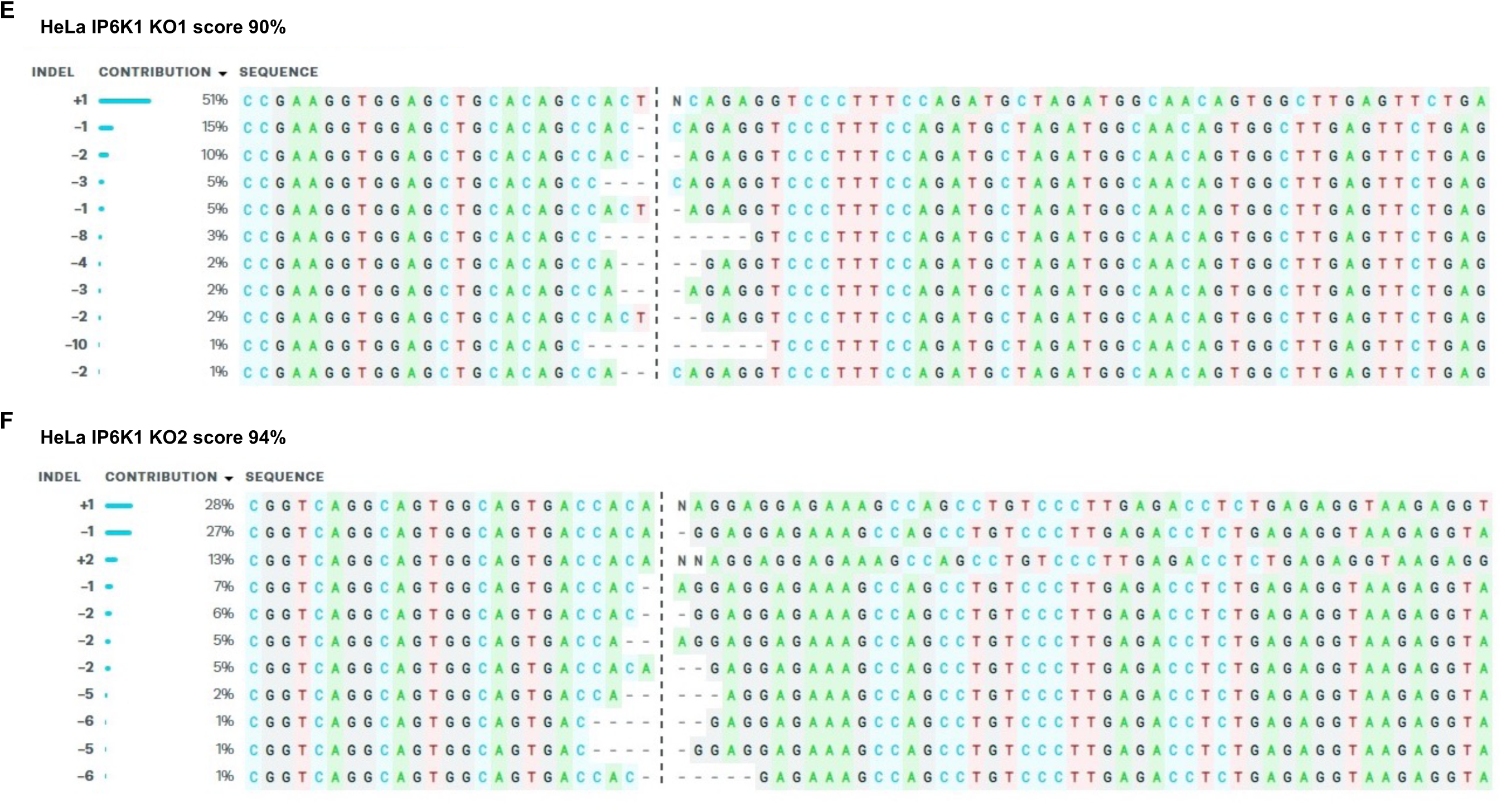
Knock out efficiency across various genetic backgrounds. (A) HEK293T ATE1 KO. (B) HEK293T ADO KO. (C) HEK293T CDO1 KO. (D) HEK293T ADO/CDO1 KO, KO efficiency is presented for CDO1, while ADO efficiency is as in (B). (E) HeLa IP6K1 KO1, (F) HeLa IP6K1 KO2. KO score calculated by ICE CRISPR analysis tool.

**Supplemental Figure 2.**
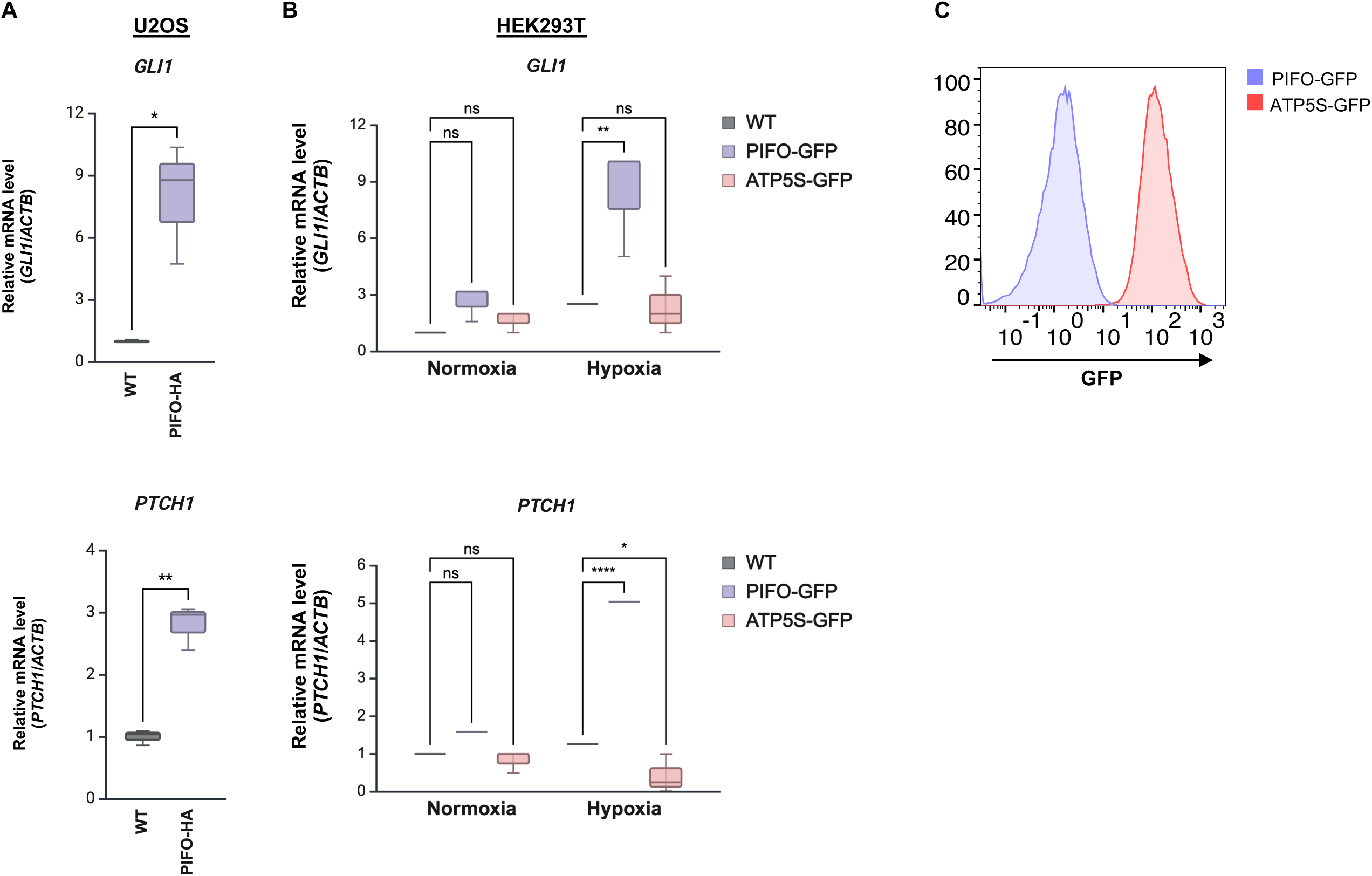
Exogenous expression of PIFO induces expression of Hedgehog signaling target genes. (A) Quantitation of the mRNA expression Hedgehog signaling pathway target genes (*GLI1*, *PTCH1*) in U2OS cells with ectopic expression of PIFO-HA. (B) Comparison of mRNA levels of Hedgehog signaling pathway genes in wild-type HEK293T cells and those expressing PIFO-GFP or ATP5S-GFP under normoxic and hypoxic conditions. ATP5S serves as an exogenous control expressed protein, as it does not influence the expression of Hedgehog signaling genes. (C) Expression levels of the GFP-fusion proteins measured using flow cytometry. Data presented as mean. Ns-no significance, P > 0.05; *P < 0.05; **P < 0.01; ***P < 0.001; ****P < 0.0001.

**Supplemental Figure S3.**
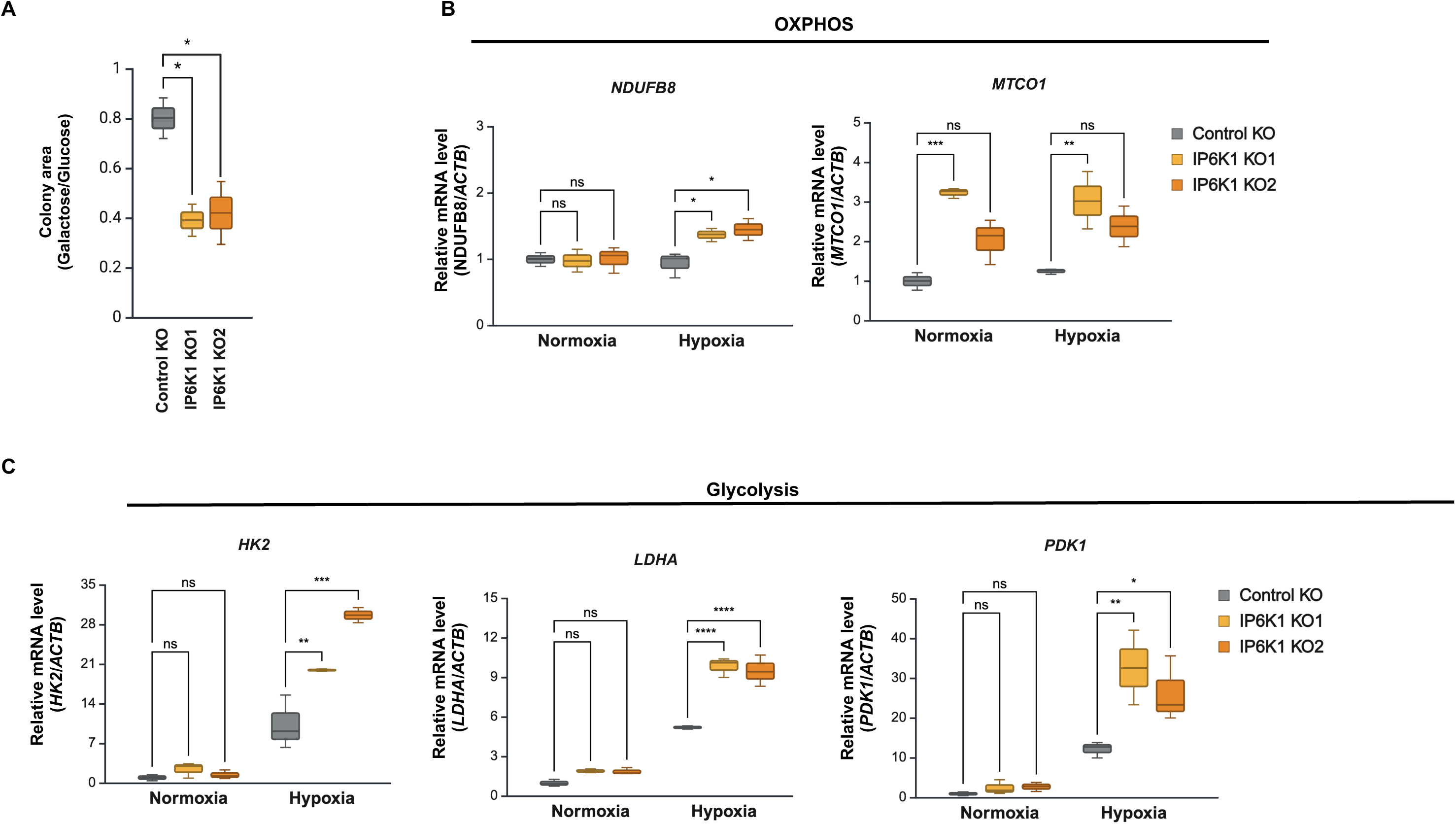
IP6K1 deficiency effect on mitochondrial metabolism and metabolic gene expression. (A) Colony formation assay of HeLa control and IP6K1 KO cells grown in media supplemented with high glucose or galactose (10 mM) for 12 days. (B-C) mRNA level normalized to *ACTB* of OXPHOS (B) or glycolytic genes (C) in control and IP6K1 KO cells cultured in normoxic versus hypoxic conditions. Data presented as mean. Ns-no significance, P > 0.05; *P < 0.05; **P < 0.01; ***P < 0.001; ****P < 0.0001.

---

**Supplemental Table 1. Candidate ATE1 substrates recovered from Ub-GPS N24mer library screen data.** (A) N24mer library. (B) N24mer MC-containing ATE1 substrates, (C) N24mer non-MC-initiating ATE1 substrates.

**Supplemental Table 2. Mutagenesis screens data.** (A) Scanning mutagenesis experiment. (B) Saturation mutagenesis experiment.

## Notes

### Competing Interest Statement

The authors have declared no competing interest.

